# Antigenic characterization of the SARS-CoV-2 Omicron subvariant BA.2.75

**DOI:** 10.1101/2022.07.31.502235

**Authors:** Qian Wang, Sho Iketani, Zhiteng Li, Yicheng Guo, Andre Yanchen Yeh, Michael Liu, Jian Yu, Zizhang Sheng, Yaoxing Huang, Lihong Liu, David D. Ho

## Abstract

The SARS-CoV-2 Omicron subvariant BA.2.75 emerged recently and appears to be spreading rapidly. It has nine mutations in its spike compared to BA.2, raising concerns it may further evade vaccine-elicited and therapeutic antibodies. Here, we found BA.2.75 to be moderately more neutralization resistant to sera from vaccinated/boosted individuals than BA.2 (1.8-fold), similar to BA.2.12.1 (1.1-fold), but more neutralization sensitive than BA.4/5 (0.6-fold). Relative to BA.2, BA.2.75 showed heightened resistance to class 1 and class 3 monoclonal antibodies to the receptor-binding domain, while gaining sensitivity to class 2 antibodies. The resistance was largely conferred by the G446S and R460K mutations. Of note, BA.2.75 was slightly resistant (3.7-fold) to bebtelovimab, the only therapeutic antibody with potent activity against all Omicron subvariants. BA.2.75 also exhibited higher receptor binding affinity than other Omicron subvariants. BA.2.75 provides yet another example of the ongoing evolution of SARS-CoV-2 as it gains transmissibility while incrementally evading antibody neutralization.

## Main text

The COVID-19 pandemic is currently dominated by the SARS-CoV-2 (severe acute respiratory syndrome coronavirus 2) Omicron subvariant BA.5. Yet another subvariant known as BA.2.75 has recently emerged from India, where it has spread rapidly, displacing the predominant BA.2 subvariant locally. Moreover, BA.2.75 has now been identified in at least 25 countries worldwide^1^ (**Extended Data Fig. 1a**). Though a descendent from BA.2, it contains a distinct set of mutations in its spike protein, including five substitutions in the N-terminal domain (NTD), K147E, W152R, F157L, I210V, and G257S, and four substitutions in the receptor-binding domain (RBD), D339H, G446S, N460K, and R493Q (**Extended Data Fig. 1b**). Many of these mutations are located at sites targeted by neutralizing antibodies and may also affect the binding affinity of the spike to its receptor, angiotensin-converting enzyme 2 (ACE2). In particular, two RBD mutations, D339H and N460K, are noteworthy because they have not been identified in previous SARS-CoV-2 variants and their impacts are not yet known. Newfound variants such as BA.2.75 that are increasing in frequency raise the concern that the viruses have developed additional mechanisms to escape from neutralization by antibodies elicited by vaccination or previous infection, as well as by therapeutic monoclonal antibodies (mAbs) in clinical use. Therefore, we have evaluated the antibody evasion properties of BA.2.75, and our findings are reported here.

We first set out to profile the antigenic differences of BA.2.75 from the wildtype SARS-CoV-2 D614G and the other currently circulating Omicron subvariants BA.2, BA.2.12.1, and BA.4/5 (note that BA.4 and BA.5 share an identical spike). VSV-pseudotyped viruses of each variant were produced and then assessed for their neutralization sensitivity to sera from three different clinical cohorts: those who had received three doses of a COVID-19 mRNA vaccine (boosted) and patients with either BA.1 or BA.2 breakthrough infection after vaccination. We did not include sera from persons immunized with only two doses of COVID-19 mRNA vaccines as we had previously observed that they lacked neutralization capacity against earlier Omicron subvariants^2,3^. The clinical information for our cohorts is described in **Extended Data Table 1** and the serum neutralization profiles are shown in **Fig. 1a** and **Extended Data Fig. 2**. Consistent with previous reports^2-4^, neutralization ID_50_ (the 50% inhibitory dose) titers of the “boosted” sera were substantially lower against BA.2, BA.2.12.1, and BA.4/5 (4.9-fold, 7.8-fold, and 14.8-fold, respectively), compared with D614G. Neutralization titers against BA.2.75 were similar to those against BA.2.12.1, 8.7-fold lower than D614G, 1.8-fold lower than BA.2, but 1.7-fold higher than BA.4/5. A similar trend was observed in the “BA.1 and BA.2 breakthrough” cohorts.

**Fig. 1.**
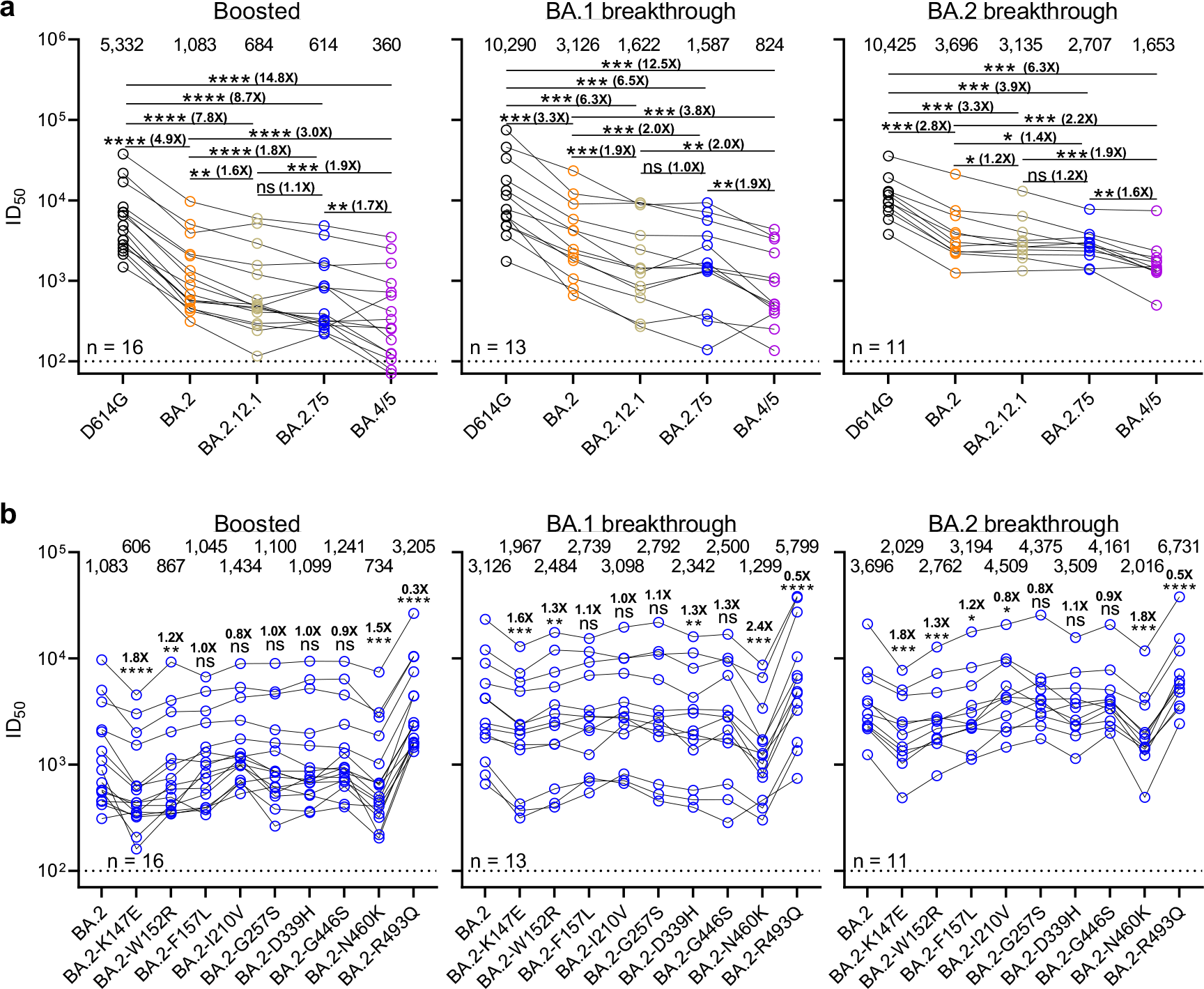
Serum neutralization profile of BA.2.75. **a**, Neutralization of pseudotyped D614G and Omicron subvariants by sera from three different clinical cohorts. Boosted refers to individuals who received three doses of a COVID-19 mRNA vaccine, and breakthrough refers to individuals who were infected and received COVID-19 vaccines. **b**, Serum neutralization of pseudotyped BA.2 or BA.2 with point mutations from BA.2.75. For both panels, values above the symbols denote the geometric mean ID_50_ values and values on the lower left indicate the sample size (n) for each group. The limit of detection (LOD) is 100 (dotted line), and values below the LOD are arbitrarily plotted to allow for visualization of each sample. *P* values were determined by using two-tailed Wilcoxon matched-pairs signed-rank tests. In **b**, comparisons were made against BA.2. Significance is denoted with asterisks and the fold-change is also denoted. ns, not significant; *, *P* < 0.05; **, *P* < 0.01; ***, *P* < 0.001; and ****, *P* < 0.0001.

We also investigated the impact of new point mutations found in BA.2.75 on serum antibody evasion by conducting serum neutralization assays with pseudoviruses containing each point mutation in the background of BA.2 (**Fig. 1b and Extended Data Fig. 3**). The mutations W152R, F157L, I210V, G257S, D339H, and N446K each only slightly (0.8-fold to 1.3-fold) altered the neutralization titers of sera from all three cohorts against BA.2. In contrast, K147E and N460K impaired the neutralization activity of sera significantly, by 1.6-fold to 1.8-fold and 1.5-fold to 2.4-fold, respectively, while the R493Q reversion mutation modestly enhanced the neutralization by 1.8-fold to 3.0-fold, as was observed previously^4^.

We next assessed the neutralization resistance of BA.2.75 to a panel of 23 mAbs directed to known neutralizing epitopes on the viral spike. Among these, 21 target the four epitope classes in the RBD^5^, including REGN10987 (imdevimab)^6^, REGN10933 (casirivimab)^6^, COV2-2196 (tixagevimab)^7^, COV2-2130 (cilgavimab)^7^, LY-CoV555 (bamlanivimab)^8^, CB6 (etesevimab)^9^, Brii-196 (amubarvimab)^10^, Brii-198 (romlusevimab)^10^, S309 (sotrovimab)^11^, LY-CoV1404 (bebtelovimab)^12^, CAB-A17^13^, ZCB11^14^, Omi-3^15^, Omi-18^15^, XGv282^16^, XGv347^16^, S2E12^17^, A19-46.1^18^, 35B5^19^, and JMB2002^20^, as well as 10-40^21^ from our group. The other two mAbs, C1717^22^ and S3H3^23^, target NTD-SD2 and SD1, respectively. Many of the recently isolated mAbs were chosen because they could neutralize earlier Omicron subvariants^13-20,22,23^.

The results are presented in **Fig. 2a** as well as in **Extended Data Fig. 4** and **Extended Data Table 2**. Unlike BA.2.12.1, which was largely similar in its antigenic profile to BA.2, BA.2.75 differed from BA.2 across a wide range of the mAbs tested. Antibodies across multiple classes were impaired against BA.2.75 compared to BA.2, including some in class 1 (CAB-A17, Omi-3, and Omi-18) and class 3 (XGv282, LY-CoV1404, JMB2002, COV2-2130, and REGN10987). Simultaneously, several antibodies showed neutralization sensitivity against BA.2.75 relative to BA.2, all within class 2 (S2E12, COV2-2196, and REGN10933). This also contrasts that of BA.4/5, which demonstrated heightened resistance to class 2 and class 3 antibodies over BA.2, but did not regain sensitivity to any antibodies. Moreover, even within the resistance to class 3 antibodies, BA.2.75 and BA.4/5 only overlapped with additional resistance towards one antibody, JMB2002. The other impaired antibodies differed, with 35B5 and Brii-198 losing further activity against BA.4/5 over BA.2. These data suggest that BA.2.75 has evolved to extend resistance towards some class 1- and class 3-directed antibodies over BA.2, yet has regained sensitivity for a subset of class 2 antibodies. Such differences may help to interpret the observations made on polyclonal serum neutralization (**Fig. 1**).

**Fig. 2.**
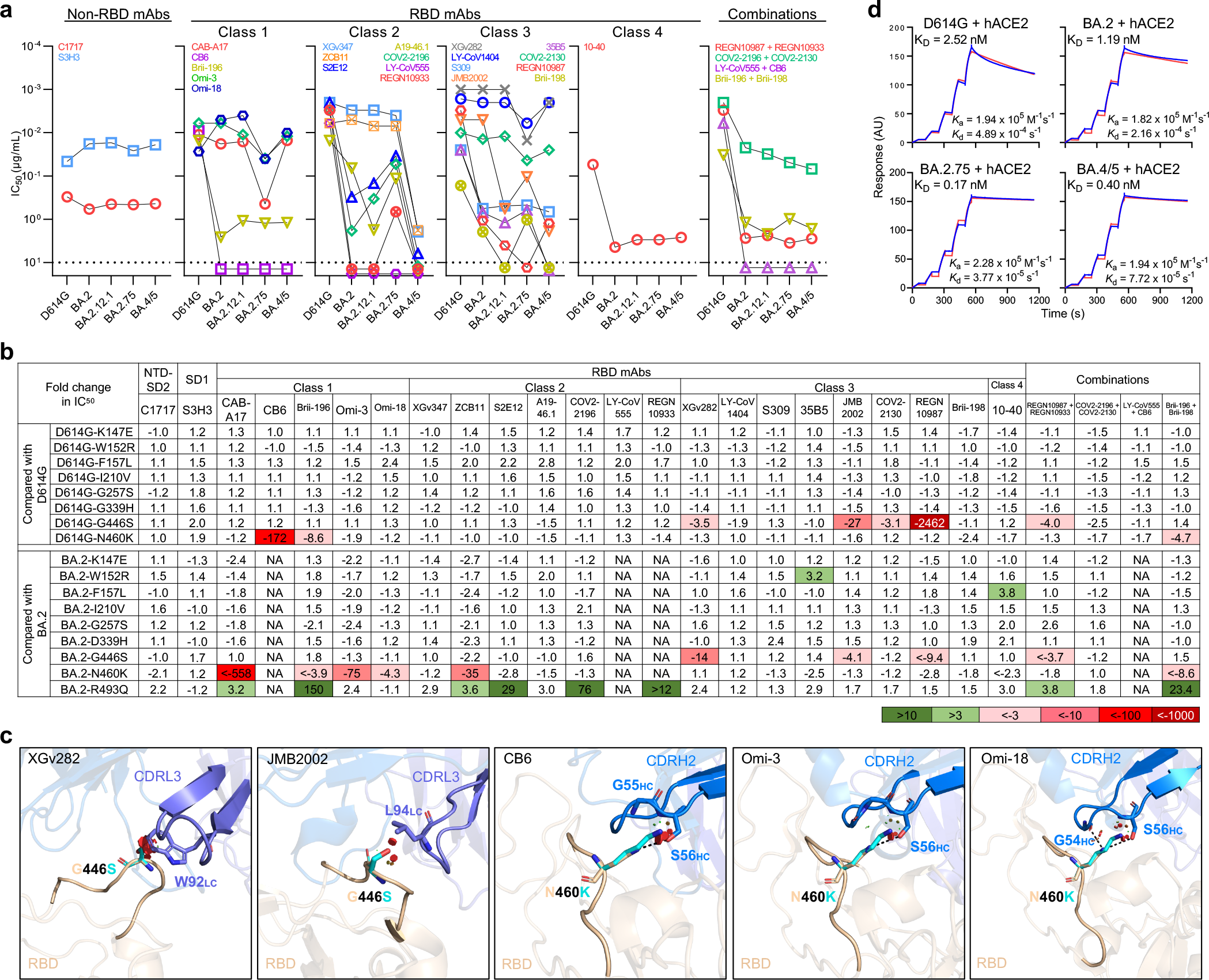
Neutralization of BA.2.75 by monoclonal antibodies and receptor binding affinity. **a**, Neutralization of pseudotyped D614G and Omicron subvariants by NTD-SD2-, SD1-, and RBD-directed mAbs. Values above the LOD of 10□μg/mL (dotted line) are arbitrarily plotted to allow for visualization of each sample. **b**, Fold change in IC_50_ values for the neutralization of pseudotyped point mutants, relative to D614G or BA.2, with resistance colored red and sensitization colored green. **c**, Modeling of the impact of G446S and N460K on antibody neutralization. Clashes are shown as red discs and hydrogen bonds are shown as dashed lines. **d**, Binding affinity of D614G, BA.2, BA.2.75, and BA.4/5 stabilized spike trimers to dimeric human ACE2. NA, not applicable.

Of particular note, BA.2.75 is the first SARS-CoV-2 variant that has demonstrated resistance to bebtelovimab (LY-CoV1404), albeit modestly at 3.7-fold loss in neutralization (**Fig. 2a**). Nevertheless, it remained the only clinical mAb that retained potent neutralizing activity against all the Omicron subvariants with a IC_50_ (the 50% inhibitory concentration) below 0.01 µg/ml. All other clinically authorized or approved antibodies or antibody combinations showed a substantial loss of activity *in vitro* against BA.2.75.

As we observed that BA.2.75 was resistant to mAbs in a unique way, we set out to identify the mutations within BA.2.75 that conferred the observed antibody resistance profile. We generated pseudoviruses carrying each of the point mutations in the background of D614G or BA.2 and tested their neutralization sensitivity to the aforementioned panel of mAbs and combinations. These data are shown in **Fig. 2b** and **Extended Data Figs. 5 and 6**. G446S impaired or abolished the neutralizing activity of class 3 mAbs (XGv282, JMB2002, and REGN10987), as previously observed in our BA.1 studies^2^. That this mutation did not result in a significant loss in polyclonal serum neutralization (**Fig. 1b**) suggests that such antibodies may be rare in a polyclonal response. The N460K substitution conferred resistance to all of the class 1 RBD mAbs tested, as well as one class 2 mAb (ZCB11). However, this resistance was only observed in the context of BA.2 for three of the class 1 antibodies (CAB-A17, Omi-3, and Omi-18) and the class 2 antibody ZCB11, but not in the context of D614G. In contrast, R493Q, also found in BA.4/5^4^, sensitized BA.2 to neutralization by several class 1 and 2 RBD mAbs, which is consistent with our previous study^4^. We note that while the NTD mutation K147E had a significant impact on polyclonal sera (**Fig. 1b**), we did not observe an effect by this mutation against the panel of mAbs tested here, suggesting that this mutation may be acting through non-RBD antibodies.

We conducted structural modeling to further investigate the impact of the G446S and N460K mutations (**Fig. 2c**). Analysis on G446S revealed steric hindrance to binding by class 3 RBD mAbs (XGv282 and JMB2002), as we reported previously^2^. In addition, structural modeling of N460K revealed that K460 abolished a common hydrogen bond between RBD-N460 and S56 in CDRH2 of VH3-53 class antibodies^5^, such as CB6, Omi-3, and Omi-18, although it is not immediately clear why the loss in activity for Omi-3 and Omi-18 was only apparent in the background of BA.2 but not D614G.

Finally, as receptor binding affinity may play a role in transmissibility, we investigated this property for BA.2.75. The binding affinity of purified spike trimer proteins of D614G, BA.2, BA.4/5, and BA.2.75 to dimeric human ACE2 (hACE2) was quantified using surface plasmon resonance (SPR). We found BA.2.75 exhibited the highest receptor binding affinity, with a K_D_ value 7.0-fold and 2.4-fold lower than values for BA.2 and BA.4/5, respectively (**Fig. 2d**). To validate these results, we tested pseudoviruses bearing these spikes for neutralization by dimeric hACE2 (**Extended Data Fig. 7**). A comparison of IC_50_ values suggested that BA.2.75 was slightly more sensitive to hACE2 than the other pseudoviruses tested, in line with was observed in the SPR. To probe the role of the mutations in BA.2.75 for ACE2 binding, we tested the neutralization by hACE2 of each of the point mutants in the background of BA.2. R493Q was the most sensitive to hACE2 neutralization, and N460K was most resistant. These results parallel our previous studies, in which we found R493Q could serve to restore the lost receptor binding affinity due to a resistance-conferring mutation in BA.4/5^4^. A similar mechanism, in which R493Q acts to balance the compromised affinity caused by N460K, may be in action for BA.2.75. In summary, we have systematically evaluated the antigenic properties of the new SARS-CoV-2 Omicron subvariant BA.2.75, which is spreading throughout the world. Our data suggest that BA.2.75 exhibits a higher resistance to vaccine-induced and infection-induced serum neutralizing activity than BA.2 (**Fig. 1a**). It is reassuring that BA.2.75 does not show greater immune evasion from polyclonal sera than the BA.4/5 subvariant (**Fig. 1a**). The resistance profile of BA.2.75 to sera can be largely attributed to the K147E and N460K mutations (**Fig. 1b**). The latter mutation is consistent with findings from deep mutational scanning of the RBD^24^. The impact of the former mutation is puzzling in that previous Omicron subvariants have already abolished the activity of the antibodies directed to the NTD antigenic supersite^2-4^ and yet this new variant evolved to contain five additional NTD mutations. Why is SARS-CoV-2 doing so when NTD antibodies contribute only a small portion of the serum virus-neutralizing activity^25,26^?

BA.2.75 exhibits a unique neutralizing profile for mAbs, with heightened resistance over BA.2 to class 1 and class 3 RBD antibodies, while gaining sensitivity towards class 2 RBD antibodies (**Fig. 2a**). Although the impairment is slight, BA.2.75 is the first SARS-CoV-2 variant to show discernible resistance to betelovimab (LY-CoV1404). Profiling of the individual mutations within BA.2.75 revealed that G446S and N460K could contribute to the resistance (**Fig. 2b**). These mutations appear to be acting through steric hindrance or abrogation of a hydrogen bond (**Fig. 2c**). More importantly, these findings on BA.2.75 demonstrate that our one remaining therapeutic antibody with potent activity to treat COVID-19 is now under threat. Another mutation proximal to residue 446 of the spike could knock out the current arsenal of therapeutic monoclonals. Although numerous mAbs have been isolated and shown to neutralize the new Omicron subvariants^13-20,22,23^, their development pathway to prove clinical efficacy has become rather daunting with the availability of effective vaccines and antiviral drugs.

Finally, our data demonstrate that BA.2.75 has enhanced binding affinity to its receptor ACE2, which may enhance its transmission (**Fig. 2d**). However, it is still unclear whether BA.2.75 could out-compete BA.5, today’s dominant form globally. The features of BA.2.75 that diverged from other Omicron subvariants serve to underscore the ability of SARS-CoV-2 to evolve, incrementally gaining transmissibility and antibody evasion, and to reinforce the importance of vaccination and booster campaigns as well as epidemiologic surveillance to detect the emergence of new SARS-CoV-2 variants.

## Methods

### Serum samples

Sera from individuals who received three doses of the mRNA-1273 or BNT162b2 vaccines were collected at Columbia University Irving Medical Center (referred to as “boosted” in the text). Sera from individuals who were infected by an Omicron subvariant (BA.1 or BA.2) following vaccinations were collected from December 2021 to May 2022 at Columbia University Irving Medical Center (referred to as “BA.1 or BA.2 breakthrough” in the text). All samples were confirmed for prior SARS-CoV-2 infection status by anti-nucleoprotein (NP) ELISA, and the variant involved in breakthrough cases were determined by sequencing. All collections were conducted under protocols reviewed and approved by the Institutional Review Board of Columbia University. All participants provided written informed consent. Clinical information on the different cohorts of study subjects is provided in **Extended Data Table 1**.

### Cell lines

Expi293 cells were obtained from Thermo Fisher Scientific (A14527); Vero-E6 cells were obtained from the ATCC (CRL-1586); HEK293T cells were obtained from the ATCC (CRL-3216). Cells were purchased from authenticated vendors and morphology was confirmed visually before use. All cell lines tested mycoplasma negative.

### Monoclonal antibodies

Antibodies were expressed in-house as previously described^27^. For each of the antibodies in this study, heavy chain variable (VH) and light chain variable (VL) genes were synthesized (GenScript), cloned into an expression vector, transfected into Expi293 cells (Thermo Fisher Scientific), and purified from the cellular supernatant by affinity purification using rProtein A Sepharose (GE). REGN10987, REGN10933, COV2-2196, and COV2-2130 were provided by Regeneron Pharmaceuticals; Brii-196 and Brii-198 were provided by Brii Biosciences; CB6 was provided by B. Zhang and P. Kwong (NIH); and ZCB11 was provided by Z. Chen (HKU).

### Construction of SARS-CoV-2 spike plasmids

Spike expression constructs for D614G, BA.2, BA.2.12.1, and BA.4/5 were previously generated^2-4^. Expression constructs encoding the BA.2.75 spike, as well as the individual mutations found in BA.2.75, were generated using the QuikChange II XL site-directed mutagenesis kit according to the manufacturer’s instructions (Agilent). For expression of stabilized soluble S2P spike trimer proteins, 2P substitutions (K986P and V987P in WA1) and a “GSAS” substitution of the furin cleavage site (682-685aa in WA1) were introduced into the spike-expressing plasmids for stabilization as previously described^28^, and then the ectodomain (1-1208aa in WA1) of the spike was fused with a C-terminal 8x His-tag and cloned into the paH vector. All constructs were confirmed by Sanger sequencing prior to use.

### Expression and purification of SARS-CoV-2 stabilized spike trimers and human ACE2

Stabilized SARS-CoV-2 spike trimer proteins of D614G and the Omicron subvariants were generated by transfecting Expi293 cells with each of the stabilized spike trimer expression constructs using 1 mg mL^-1^ polyethylenimine (PEI), and then purifying the spike trimer from the supernatants five days post-transfection using Ni-NTA resin (Invitrogen) according to the manufacturer’s instructions^27^. Dimeric human ACE2-IgG1 (hACE2) was generated by transfecting Expi293 cells with pcDNA3-sACE2-WT(732)-IgG1^29^ (Addgene plasmid #154104, gift from Erik Procko) using 1 mg mL^-1^ PEI and then purifying from the supernatant five days post-transfection using rProtein A Sepharose (GE) according to the manufacturer’s instructions. All proteins were confirmed for purity and size by SDS-PAGE prior to use.

### Surface plasmon resonance

Surface plasmon resonance (SPR) binding assays for hACE2 binding to SARS-CoV-2 stabilized spike trimers were performed using a Biacore T200 biosensor equipped with a Series S CM5 chip (Cytiva), in a running buffer of 10 mM HEPES pH 7.4, 150 mM NaCl, 3 mM EDTA, 0.05% P-20 (Cytiva) at 25 °C. Spike proteins were captured through their C-terminal His-tag over an anti-His antibody surface. These surfaces were generated using the His-capture kit (Cytiva) according to the manufacturer’s instructions, resulting in approximately 10,000 RU of anti-His antibody over each surface. An anti-His antibody surface without antigen was used as a reference flow cell to remove bulk shift changes from the binding signal. For each spike, binding to hACE2 was tested using a three-fold dilution series with concentrations ranging from 2.46 nM to 200 nM. The association and dissociation rates were each monitored for 60 s and 600 s respectively, at 30 µL/min. The bound spike/hACE2 complex was regenerated from the anti-His antibody surface using 10 mM glycine pH 1.5. Blank buffer cycles were performed by injecting running buffer instead of hACE2 remove systematic noise from the binding signal. The resulting data was processed and fit to a 1:1 binding model using Biacore Evaluation Software.

### Pseudovirus production

Pseudotyped SARS-CoV-2 (pseudoviruses) were produced in the vesicular stomatitis virus (VSV) background, in which the native VSV glycoprotein was replaced by that of SARS-CoV-2 and its variants, as previously described^27^. HEK293T cells were transfected with the appropriate spike expression construct using 1 mg mL^-1^ PEI and cultured overnight at 37□°C under 5% CO_2_, and then infected with VSV-G pseudotyped ΔG-luciferase (G*ΔG-luciferase, Kerafast) 24 h post-transfection. After 2□h of infection, cells were washed three times, changed to fresh medium, and then cultured for approximately another 24□h before the supernatants were collected, clarified by centrifugation, and aliquoted and stored at -80 °C until further use.

### Pseudovirus neutralization assay

Prior to use in the neutralization assay, all pseudoviruses were first titrated to equilibrate the viral input between assays. Five-fold serial dilutions of heat-inactivated sera, antibodies, or hACE2 were prepared in media in 96-well plates in triplicate, starting at 1:100 dilution for sera and 10□µg□mL^−1^ for antibodies and hACE2. Pseudoviruses were then added to wells and the virus– sample mixture was incubated at 37□°C for 1□h, except for hACE2, where no incubation was conducted. Control wells with the virus only were included on all plates. Vero-E6 cells were then added at a density of 3□×□10^4^ cells per well and the plates were incubated at 37□°C for approximately 10□h. Cells were then lysed and luciferase activity was quantified using the Luciferase Assay System (Promega) according to the manufacturer’s instructions with SoftMax Pro v.7.0.2 (Molecular Devices). Neutralization curves and IC_50_ values were derived by fitting a nonlinear five-parameter dose-response curve to the data in GraphPad Prism v.9.2.

### Structural modeling of RBD mutations

The structures of antibody-spike complexes for modeling were obtained from PDB (7WLC (XGv282), 7XOD (JMB2002), 7C01 (CB6), 7ZF3 (Omi-3), and 7ZFB (Omi-18)). PyMOL v.2.3.2 was used to perform mutagenesis, to identify steric clashes and hydrogen bonds between RBD and antibodies, and to generate structural plots (Schrödinger, LLC).

## Acknowledgements

This study was supported by funding from the Gates Foundation, JPB Foundation, Andrew and Peggy Cherng, Samuel Yin, Carol Ludwig, David and Roger Wu, Regeneron Pharmaceuticals, and the NIH SARS-CoV-2 Assessment of Viral Evolution (SAVE) Program. We are grateful to Michael T. Yin, Magdalena E. Sobieszczyk, Jennifer Y. Chang, Jayesh G. Shah, and David S. Perlin for providing serum samples from COVID-19 patients. We thank all who have contributed their data to GISAID.

## Author contributions

D.D.H. and L.L. conceived this project. Q.W., A.Y.Y., and L.L. constructed the spike expression plasmids and produced pseudoviruses. Q.W., S.I., and L.L. conducted pseudovirus neutralization experiments. Q.W., Z.L., A.Y.Y., and L.L. purified SARS-CoV-2 spike and ACE2 proteins and performed SPR. Y.G. and Z.S. conducted bioinformatic analyses. Q.W. managed the project. J.Y. and M.L. produced antibodies. D.D.H. and L.L. directed and supervised the project. Q.W., S.I., Y.G., L.L., and D.D.H. analyzed the results and wrote the manuscript.

## Additional information

Correspondence and requests for materials should be addressed to L.L. or D.D.H.

## Data availability

All data are provided in the manuscript. Materials in this study will be made available under an appropriate Materials Transfer Agreement. Sequences for BA.2.75 prevalence analysis were downloaded from GISAID (https://www.gisaid.org/). The structures used for analysis in this study are available from PDB under IDs 7WLC (XGv282), 7XOD (JMB2002), 7C01 (CB6), 7ZF3 (Omi-3), and 7ZFB (Omi-18).

## Extended Data Legends

**Extended Data Fig. 1** | **Prevalence of BA.2.75 and mutations found in BA.2.75 spike. a**, The cumulative number of BA.2.75 sequences found in India and globally, and the number of countries reporting BA.2.75 cases, were tabulated from GISAID. **b**, Spike mutations found in BA.2.75 relative to BA.2.

**Extended Data Fig. 2** | **Neutralization curves for serum against pseudotyped D614G and Omicron subvariants**. Neutralization by **a**, boosted vaccinee sera on pseudoviruses. **b**, BA.1 breakthrough sera on pseudoviruses. **c**, BA.2 breakthrough sera on pseudoviruses. Error bars denote mean ± SEM for three technical replicates.

**Extended Data Fig. 3** | **Neutralization curves for serum against pseudotyped BA.2 and BA.2 carrying individual mutations from BA.2.75**. Neutralization by **a**, boosted vaccinee sera. **b**, BA.1 breakthrough sera. **c**, BA.2 breakthrough sera. Error bars denote mean ± SEM for three technical replicates.

**Extended Data Fig. 4** | **Neutralization curves for mAbs against pseudotyped D614G and Omicron subvariants**. Data are shown as mean ± SEM from three technical replicates.

**Extended Data Fig. 5** | **Neutralization curves for mAbs against pseudotyped D614G carrying individual mutations from BA.2.75**. Data are shown as mean ± SEM from three technical replicates.

**Extended Data Fig. 6** | **Neutralization curves for mAbs against pseudotyped BA.2 carrying individual mutations from BA.2.75**. Data are shown as mean ± SEM from three technical replicates.

**Extended Data Fig. 7** | **Neutralization of pseudotyped D614G and Omicron subvariants by ACE2**. Sensitivity of pseudotyped D614G, Omicron subvariants, and BA.2 carrying individual mutations from BA.2.75 to hACE2 inhibition. The hACE2 concentrations resulting in 50% inhibition of infectivity (IC_50_) are denoted. Data are shown as mean ± standard error of mean (SEM) for three technical replicates.

**Extended Data Table 1** | **Demographics of clinical cohorts in this study**.

**Extended Data Table 2** | **IC**_**50**_ **values for pseudovirus neutralization by mAbs against D614G, Omicron subvariants, and D614G and BA.2 carrying individual mutations from BA.2.75**.

