## Supplemental figures for "Antigenic characterization of the SARS-CoV-2 Omicron subvariant BA.2.75"

### Extended Data Fig. 1

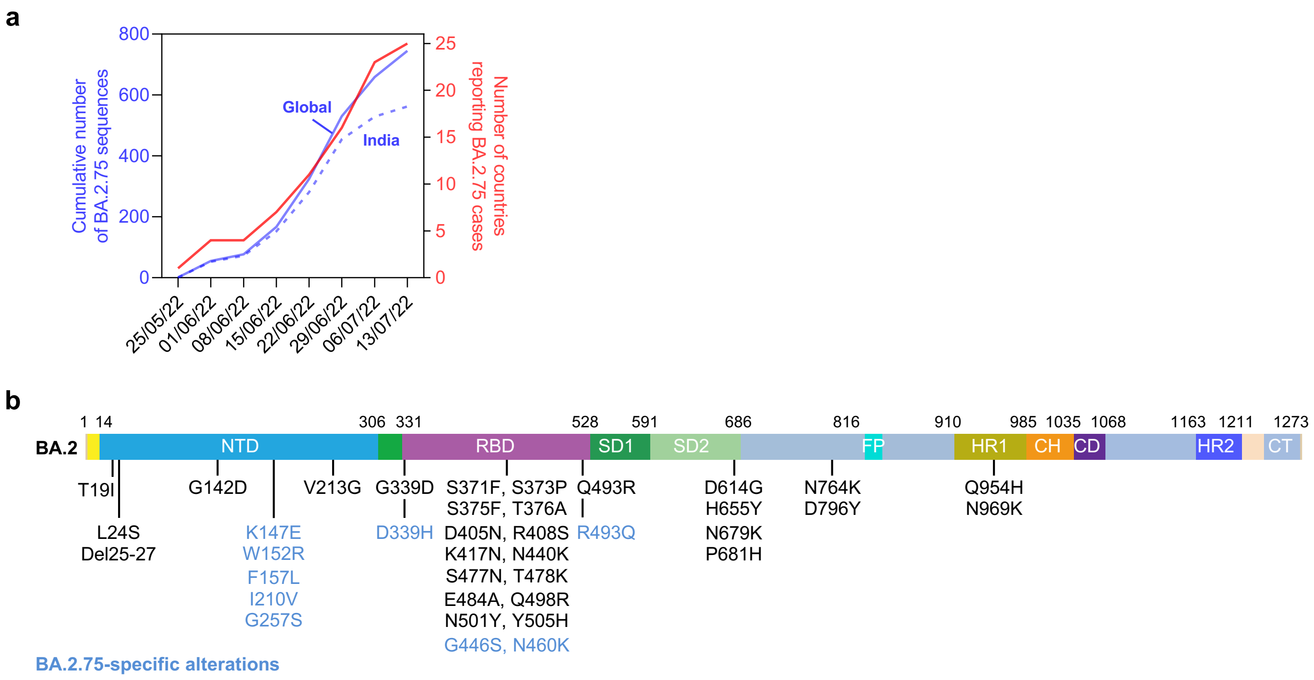

### Extended Data Fig. 2

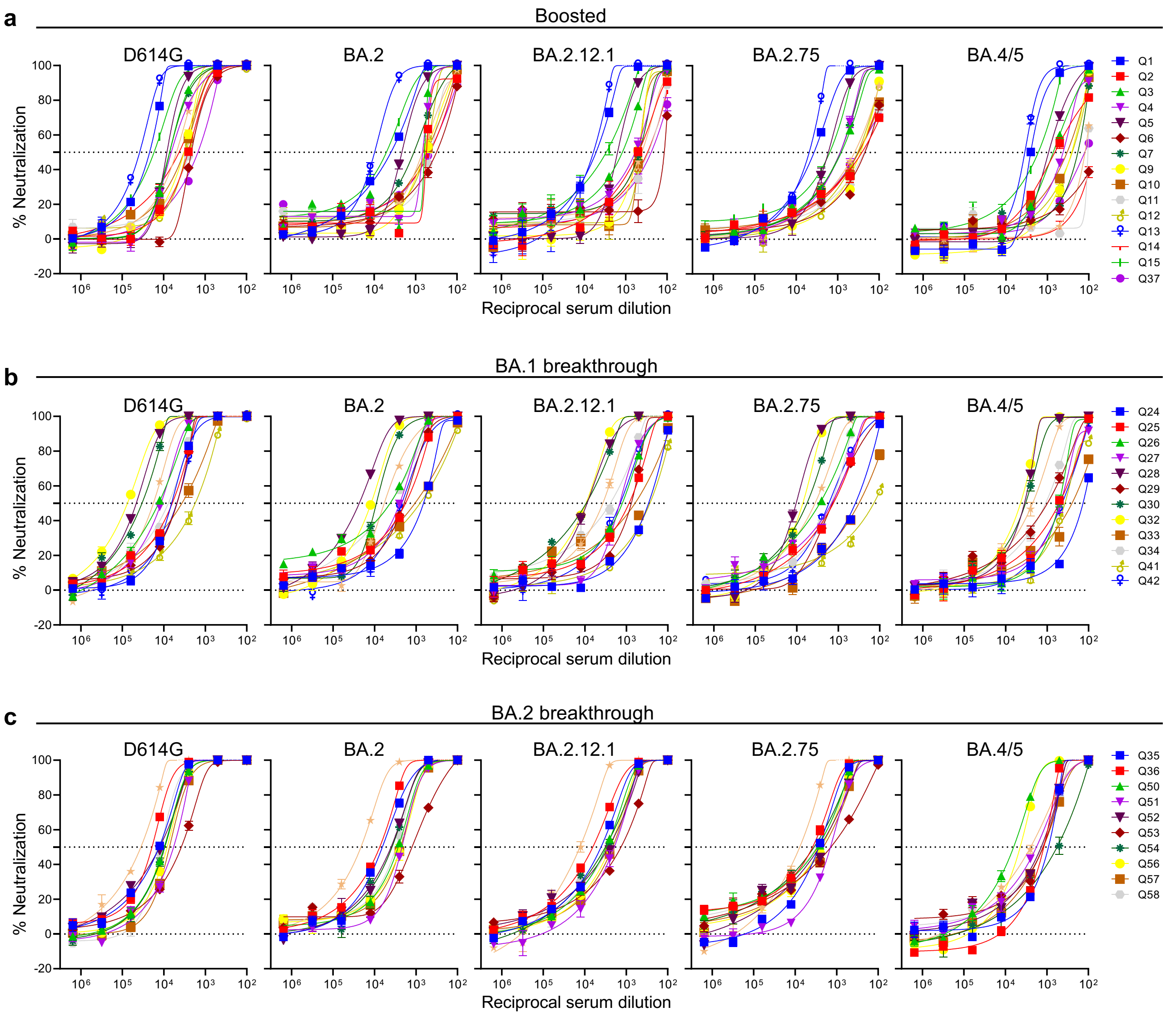

### Extended Data Fig. 3

**a**

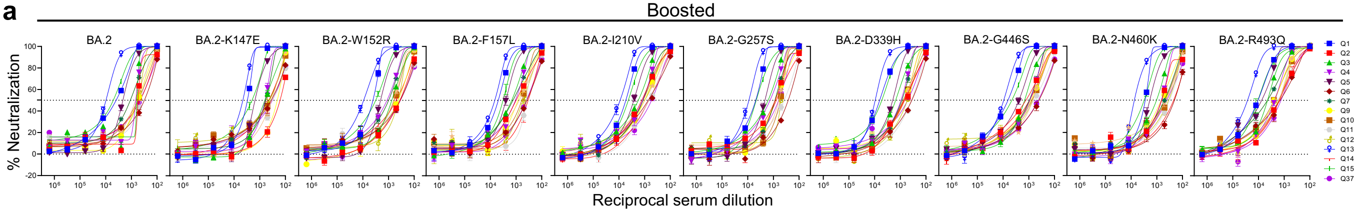


**b**

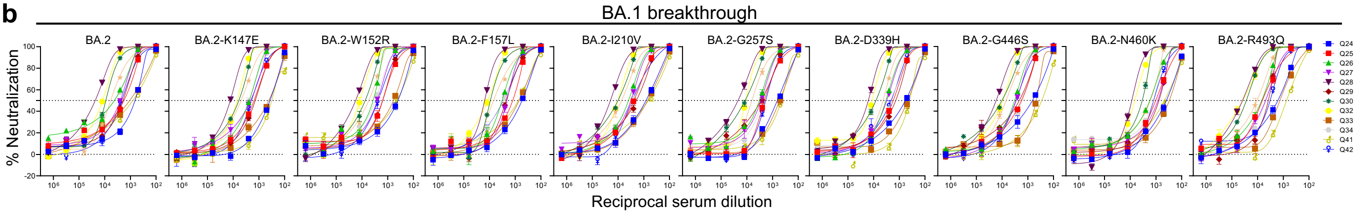


**c**

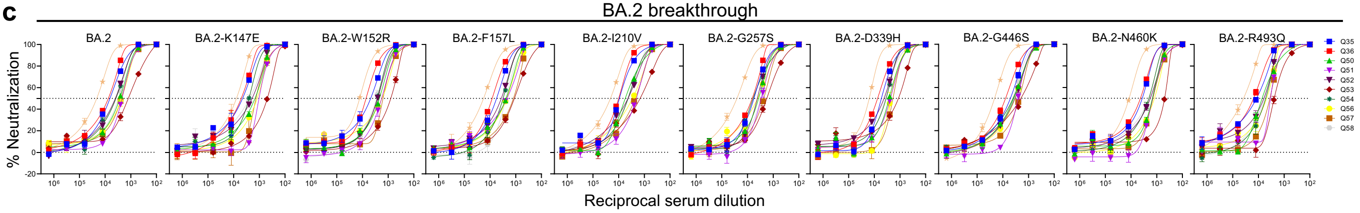

### Extended Data Fig. 4

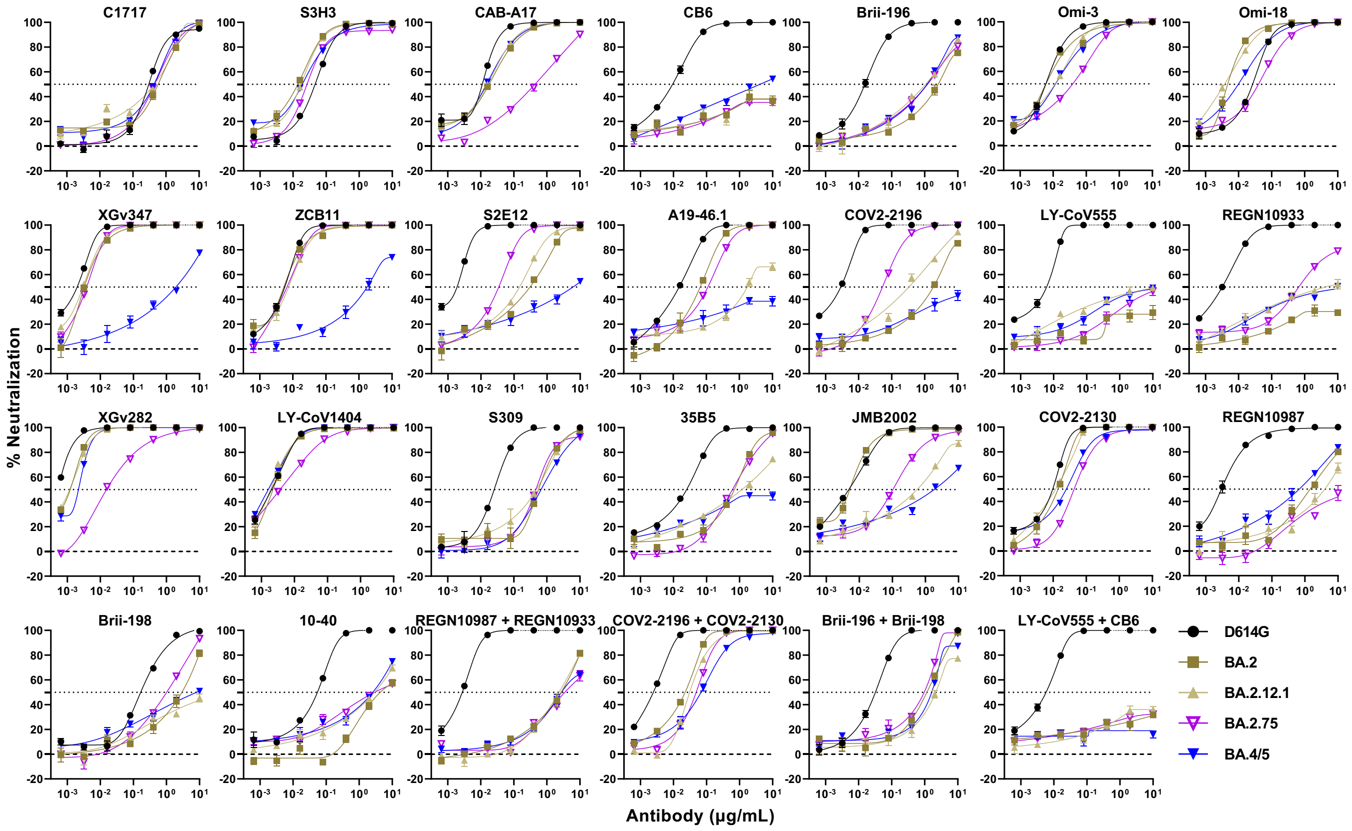

### Extended Data Fig. 5

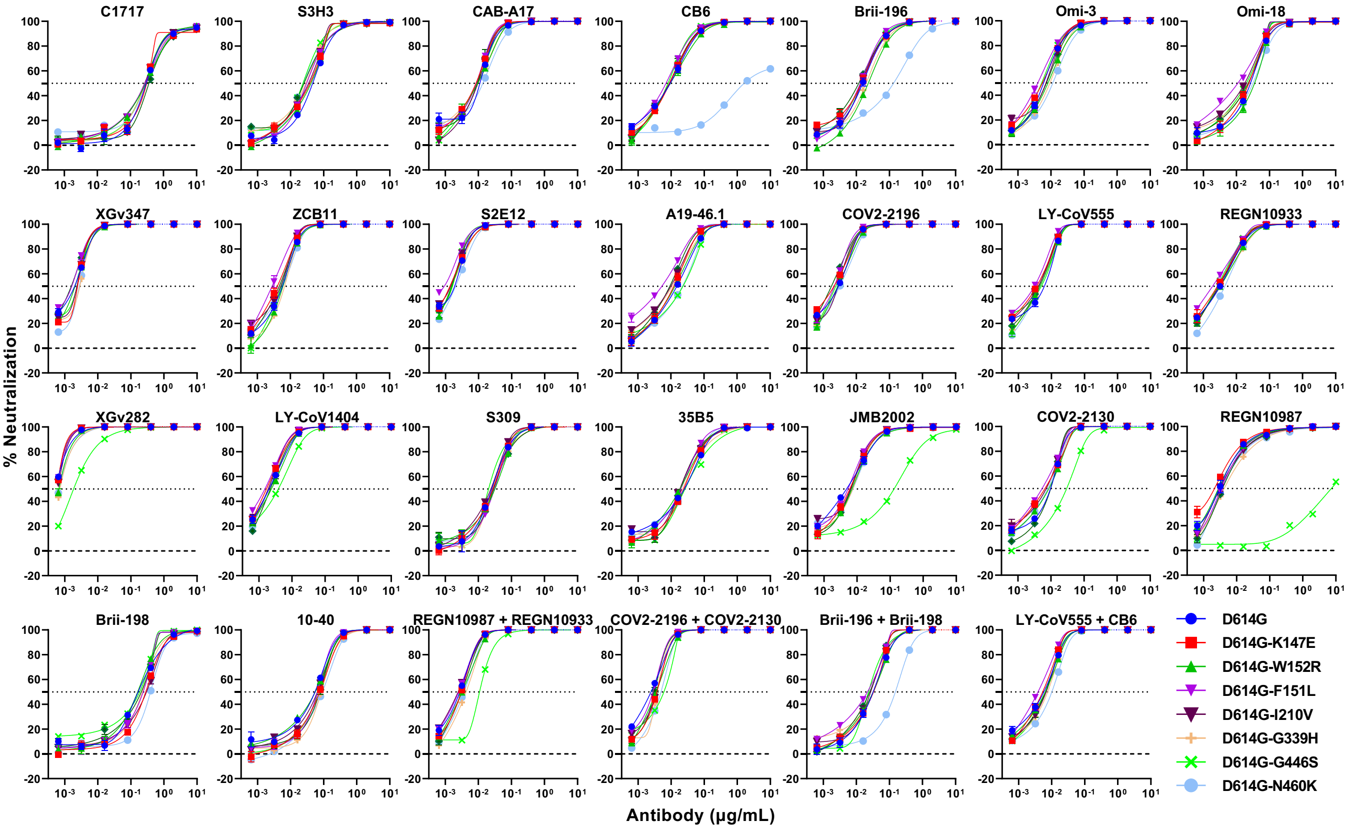

### Extended Data Fig. 6

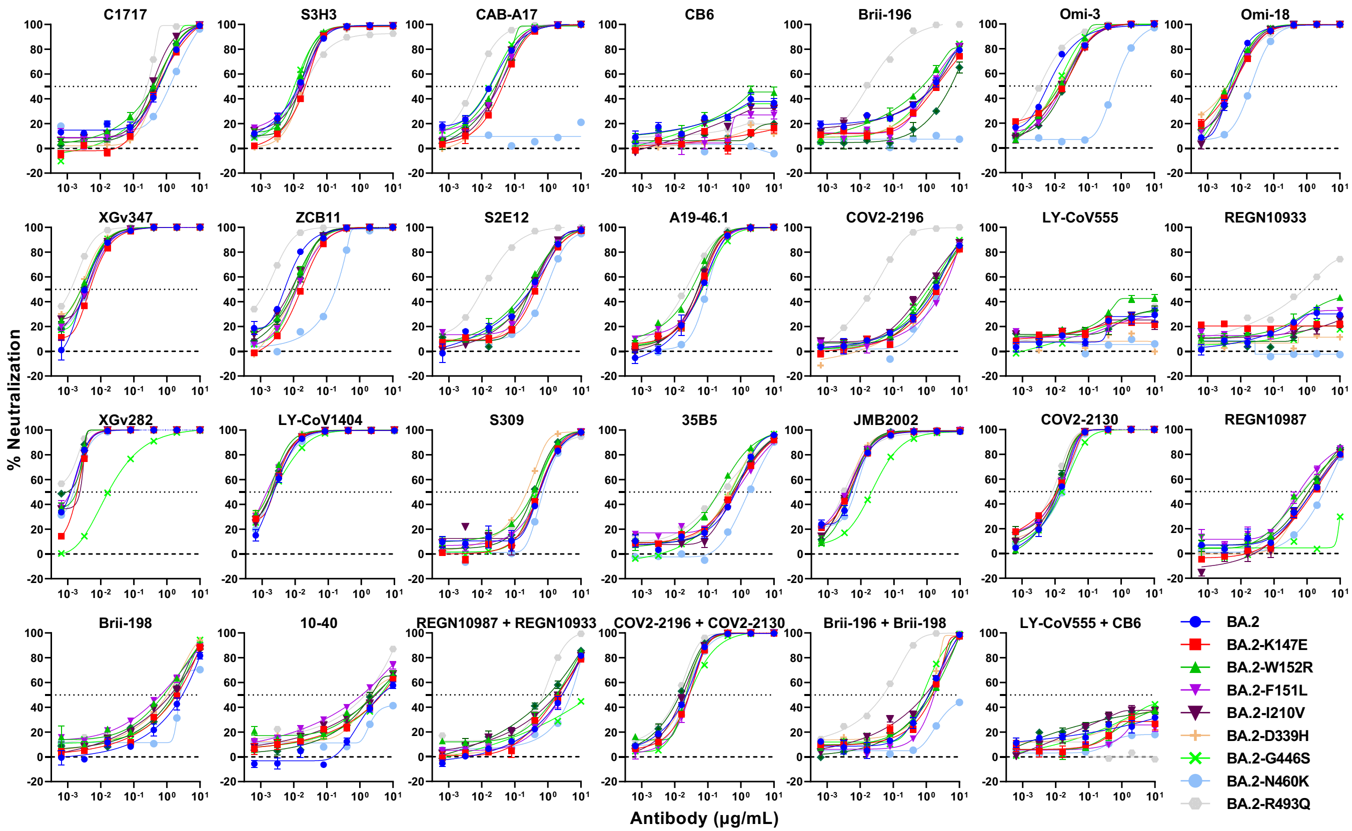

### Extended Data Fig. 7

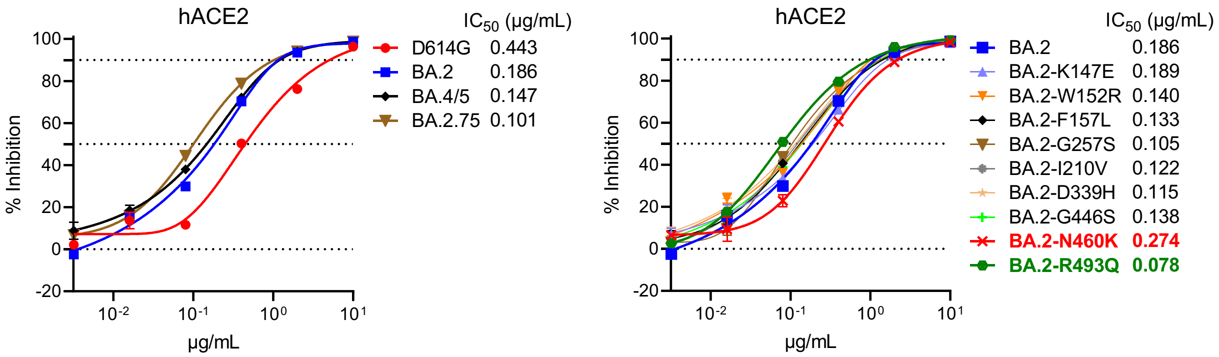

### Extended Data Table 1

| Sample ID | Vaccine type and infected strain | Days post-vaccination or *infection (after last exposure) | Documented COVID-19 | Age | Gender |
|---|---|---|---|---|---|
| Boosted | | | | | |
| Q1 | mRNA-1273/mRNA-1273/mRNA-1273 | 29 | No | 66 | Female |
| Q2 | BNT162b2/BNT162b2/BNT162b2 | 30 | No | 68 | Male |
| Q3 | BNT162b2/BNT162b2/BNT162b2 | 14 | No | 64 | Female |
| Q4 | BNT162b2/BNT162b2/BNT162b2 | 34 | No | 55 | Male |
| Q5 | BNT162b2/BNT162b2/BNT162b2 | 34 | No | 45 | Male |
| Q6 | BNT162b2/BNT162b2/BNT162b2 | 15 | No | 50 | Female |
| Q7 | BNT162b2/BNT162b2/BNT162b2 | 15 | No | 48 | Female |
| Q8 | BNT162b2/BNT162b2/BNT162b2 | 29 | No | 71 | Male |
| Q9 | BNT162b2/BNT162b2/BNT162b2 | 90 | No | 59 | Male |
| Q10 | BNT162b2/BNT162b2/BNT162b2 | 33 | No | 45 | Male |
| Q11 | BNT162b2/BNT162b2/BNT162b2 | 87 | No | 66 | Female |
| Q12 | BNT162b2/BNT162b2/BNT162b2 | 84 | No | 26 | Male |
| Q13 | mRNA-1273/mRNA-1273/mRNA-1273 | 23 | No | 28 | Female |
| Q14 | BNT162b2/BNT162b2/BNT162b2 | 14 | No | 78 | Male |
| Q15 | BNT162b2/BNT162b2/mRNA-1273 | 32 | No | 39 | Male |
| Q37 | BNT162b2/BNT162b2/BNT162b2 | 20 | No | Unknown | Female |
| BA.1 breakthrough | | | | | |
| Q24 | BNT162b2/BNT162b2/BA.1 | *14 | Yes | Unknown | Unknown |
| Q25 | BNT162b2/BNT162b2/BA.1 | *14 | Yes | Unknown | Unknown |
| Q26 | mRNA-1273/mRNA-1273/BA.1 | *35 | Yes | Unknown | Unknown |
| Q27 | BNT162b2/BNT162b2/BNT162b2/BA.1 | *135 | Yes | 78 | Male |
| Q28 | BNT162b2/BNT162b2/BNT162b2/BA.1 | *14 | Yes | Unknown | Unknown |
| Q29 | BNT162b2/BNT162b2/BNT162b2/BA.1 | *14 | Yes | Unknown | Unknown |
| Q30 | BNT162b2/BNT162b2/BNT162b2/BA.1 | *14 | Yes | Unknown | Unknown |
| Q31 | BNT162b2/BNT162b2/BNT162b2/BA.1 | *41 | Yes | 48 | Male |
| Q32 | BNT162b2/BNT162b2/BNT162b2/BA.1 | *26 | Yes | 38 | Female |
| Q33 | BNT162b2/BNT162b2/B.1.617.2/BNT162b2/BA.1 | *19 | Yes | 35 | Female |
| Q34 | BNT162b2/BNT162b2/mRNA-1273/mRNA-1273/BA.1 | *67 | Yes | 40 | Male |
| Q41 | WA1/BNT162b2/BA.1 | *21 | Yes | 52 | Male |
| Q42 | WA1/BNT162b2/BA.1 | *44 | Yes | 37 | Intersex |
| BA.2 breakthrough | | | | | |
| Q35 | BNT162b2/BNT162b2/BA.2 | *14 | Yes | 50 | Female |
| Q36 | BNT162b2/BNT162b2/BNT162b2/Ad26.COV2.S/BA.2 | *22 | Yes | 69 | Male |
| Q50 | mRNA-1273/mRNA-1273/mRNA-1273/BA.2 | *14 | Yes | 34 | Male |
| Q51 | BNT162b2/BNT162b2/mRNA-1273/BA.2 | *19 | Yes | 33 | Female |
| Q52 | BNT162b2/BNT162b2/mRNA-1273/BA.2 | *18 | Yes | 29 | Female |
| Q53 | BNT162b2/BNT162b2/BNT162b2/BA.2 | *25 | Yes | 34 | Male |
| Q54 | BNT162b2/BNT162b2/BNT162b2/BA.2 | *36 | Yes | 37 | Female |
| Q55 | BNT162b2/BNT162b2/mRNA-1273/BA.2 | *18 | Yes | 41 | Female |
| Q56 | mRNA-1273/mRNA-1273/mRNA-1273/BA.2 | *21 | Yes | 36 | Female |
| Q57 | BNT162b2/BNT162b2/mRNA-1273/BA.2 | *32 | Yes | 28 | Male |
| Q58 | BNT162b2/BNT162b2/mRNA-1273/BA.2 | *23 | Yes | 33 | Female |

### Extended Data Table 2

| $IC_{50}$ (µg/mL) | NTD-SD2 | SD1 | RBD | | | | | | | | | | | | | | | | | | | | | Combination | | | |
|---|---|---|---|---|---|---|---|---|---|---|---|---|---|---|---|---|---|---|---|---|---|---|---|---|---|---|---|
| | | | Class 1 | | | | | Class 2 | | | | | | | Class 3 | | | | | | | | Class 4 | | | | |
| | C1717 | S3H3 | CAB-A17 | CB6 | Brii-196 | Omi-3 | Omi-18 | XGv347 | ZCB11 | S2E12 | A19-46.1 | COV2-2196 | LY-CoV555 | REGN 10933 | XGv282 | LY-CoV1404 | S309 | 35B5 | JMB2002 | COV2-2130 | REGN 10987 | Brii-198 | 10-40 | REGN10987 + REGN10933 | COV2-2196 + COV2-2130 | LY-CoV555 + CB6 | Brii-196 + Brii-198 |
| D614G | 0.302 | 0.046 | 0.011 | 0.009 | 0.015 | 0.006 | 0.027 | 0.002 | 0.006 | 0.002 | 0.015 | 0.003 | 0.006 | 0.003 | <0.001 | 0.002 | 0.025 | 0.025 | 0.005 | 0.010 | 0.003 | 0.166 | 0.054 | 0.003 | 0.002 | 0.006 | 0.032 |
| BA.2 | 0.579 | 0.018 | 0.018 | >10 | 2.555 | 0.006 | 0.005 | 0.003 | 0.005 | 0.305 | 0.065 | 1.825 | >10 | >10 | 0.001 | 0.002 | 0.558 | 0.697 | 0.005 | 0.014 | 1.066 | 1.931 | 4.355 | 2.701 | 0.022 | >10 | 1.164 |
| BA.2.12.1 | 0.445 | 0.017 | 0.016 | >10 | 1.066 | 0.011 | 0.004 | 0.003 | 0.007 | 0.147 | 1.737 | 0.334 | >10 | >10 | 0.001 | 0.002 | 0.484 | 1.185 | 0.561 | 0.012 | 4.002 | >10 | 2.936 | 2.343 | 0.031 | >10 | 2.155 |
| BA.2.75 | 0.457 | 0.026 | 0.438 | >10 | 1.214 | 0.039 | 0.040 | 0.004 | 0.007 | 0.034 | 0.112 | 0.053 | >10 | 0.664 | 0.015 | 0.006 | 0.467 | 0.609 | 0.104 | 0.043 | >10 | 1.016 | 2.930 | 3.504 | 0.049 | >10 | 0.963 |
| BA.4/5 | 0.436 | 0.019 | 0.015 | >10 | 1.181 | 0.012 | 0.010 | 1.935 | 1.774 | 6.115 | >10 | >10 | >10 | >10 | 0.002 | 0.002 | 0.669 | >10 | 1.847 | 0.025 | 1.230 | >10 | 2.590 | 2.779 | 0.068 | >10 | 1.625 |
| D614G-K147E | 0.311 | 0.038 | 0.008 | 0.009 | 0.014 | 0.006 | 0.024 | 0.002 | 0.004 | 0.001 | 0.012 | 0.002 | 0.004 | 0.003 | <0.001 | 0.002 | 0.028 | 0.024 | 0.006 | 0.007 | 0.002 | 0.282 | 0.076 | 0.003 | 0.004 | 0.005 | 0.033 |
| D614G-W152R | 0.295 | 0.041 | 0.010 | 0.009 | 0.023 | 0.008 | 0.035 | 0.002 | 0.006 | 0.002 | 0.013 | 0.003 | 0.004 | 0.003 | <0.001 | 0.003 | 0.032 | 0.019 | 0.007 | 0.009 | 0.003 | 0.160 | 0.056 | 0.003 | 0.004 | 0.006 | 0.032 |
| D614G-F157L | 0.278 | 0.031 | 0.009 | 0.007 | 0.013 | 0.004 | 0.011 | 0.002 | 0.003 | <0.001 | 0.005 | 0.002 | 0.003 | 0.002 | <0.001 | 0.002 | 0.031 | 0.020 | 0.005 | 0.005 | 0.003 | 0.236 | 0.065 | 0.003 | 0.003 | 0.004 | 0.022 |
| D614G-I210V | 0.269 | 0.035 | 0.010 | 0.009 | 0.014 | 0.007 | 0.018 | 0.002 | 0.005 | 0.001 | 0.010 | 0.003 | 0.004 | 0.003 | <0.001 | 0.002 | 0.024 | 0.018 | 0.006 | 0.007 | 0.003 | 0.298 | 0.065 | 0.002 | 0.004 | 0.007 | 0.027 |
| D614G-G257S | 0.364 | 0.025 | 0.009 | 0.009 | 0.012 | 0.007 | 0.023 | 0.002 | 0.005 | 0.002 | 0.009 | 0.002 | 0.004 | 0.003 | <0.001 | 0.002 | 0.029 | 0.019 | 0.007 | 0.010 | 0.004 | 0.206 | 0.077 | 0.003 | 0.003 | 0.006 | 0.025 |
| D614G-G339H | 0.288 | 0.029 | 0.010 | 0.009 | 0.020 | 0.009 | 0.023 | 0.003 | 0.007 | 0.002 | 0.011 | 0.003 | 0.004 | 0.003 | <0.001 | 0.002 | 0.027 | 0.022 | 0.007 | 0.007 | 0.004 | 0.221 | 0.084 | 0.004 | 0.004 | 0.008 | 0.033 |
| D614G-G446S | 0.278 | 0.024 | 0.009 | 0.008 | 0.014 | 0.005 | 0.020 | 0.002 | 0.005 | 0.002 | 0.023 | 0.003 | 0.005 | 0.003 | 0.002 | 0.004 | 0.020 | 0.026 | 0.132 | 0.030 | 7.311 | 0.185 | 0.046 | 0.011 | 0.006 | 0.007 | 0.023 |
| D614G-N460K | 0.290 | 0.024 | 0.014 | 1.622 | 0.133 | 0.011 | 0.031 | 0.003 | 0.006 | 0.002 | 0.022 | 0.003 | 0.005 | 0.004 | <0.001 | 0.002 | 0.029 | 0.026 | 0.008 | 0.008 | 0.004 | 0.392 | 0.091 | 0.004 | 0.004 | 0.010 | 0.151 |
| BA.2-K147E | 0.533 | 0.022 | 0.042 | >10 | 1.987 | 0.018 | 0.005 | 0.005 | 0.015 | 0.428 | 0.059 | 2.174 | >10 | >10 | 0.002 | 0.002 | 0.532 | 0.606 | 0.004 | 0.012 | 1.625 | 1.973 | 4.524 | 1.993 | 0.027 | >10 | 1.523 |
| BA.2-W152R | 0.389 | 0.013 | 0.025 | >10 | 1.397 | 0.014 | 0.004 | 0.003 | 0.009 | 0.209 | 0.032 | 1.614 | >10 | >10 | 0.001 | 0.002 | 0.364 | 0.216 | 0.005 | 0.012 | 0.764 | 1.361 | 2.703 | 1.812 | 0.021 | >10 | 1.436 |
| BA.2-F157L | 0.584 | 0.016 | 0.033 | >10 | 1.319 | 0.017 | 0.006 | 0.004 | 0.013 | 0.365 | 0.063 | 3.181 | >10 | >10 | 0.001 | 0.001 | 0.563 | 0.709 | 0.004 | 0.012 | 0.582 | 1.389 | 1.144 | 2.698 | 0.028 | >10 | 1.747 |
| BA.2-I210V | 0.352 | 0.017 | 0.028 | >10 | 1.674 | 0.016 | 0.006 | 0.004 | 0.009 | 0.299 | 0.051 | 0.878 | >10 | >10 | 0.002 | 0.002 | 0.526 | 0.639 | 0.004 | 0.012 | 1.381 | 1.296 | 2.990 | 1.786 | 0.018 | >10 | 0.918 |
| BA.2-G257S | 0.474 | 0.015 | 0.033 | >10 | 5.475 | 0.020 | 0.006 | 0.003 | 0.011 | 0.299 | 0.050 | 1.419 | >10 | >10 | <0.001 | 0.002 | 0.375 | 0.562 | 0.004 | 0.010 | 1.098 | 1.531 | 2.222 | 1.033 | 0.014 | >10 | 1.218 |
| BA.2-D339H | 0.522 | 0.018 | 0.029 | >10 | 1.731 | 0.013 | 0.004 | 0.002 | 0.012 | 0.287 | 0.050 | 2.170 | >10 | >10 | 0.001 | 0.002 | 0.229 | 0.462 | 0.004 | 0.011 | 1.116 | 1.033 | 2.093 | 2.569 | 0.021 | >10 | 1.182 |
| BA.2-G446S | 0.582 | 0.010 | 0.017 | >10 | 1.408 | 0.011 | 0.005 | 0.003 | 0.012 | 0.313 | 0.069 | 1.140 | >10 | >10 | 0.017 | 0.002 | 0.450 | 0.487 | 0.021 | 0.016 | >10 | 1.778 | 4.196 | >10 | 0.027 | >10 | 0.780 |
| BA.2-N460K | 1.236 | 0.014 | >10 | >10 | >10 | 0.625 | 0.020 | 0.004 | 0.187 | 0.848 | 0.096 | 2.421 | >10 | >10 | 0.001 | 0.002 | 0.712 | 1.721 | 0.007 | 0.017 | 3.029 | 3.521 | >10 | 4.905 | 0.022 | >10 | >10 |
| BA.2-R493Q | 0.260 | 0.021 | 0.006 | >10 | 0.017 | 0.003 | 0.005 | 0.001 | 0.001 | 0.011 | 0.022 | 0.024 | >10 | 0.836 | <0.001 | 0.002 | 0.416 | 0.243 | 0.003 | 0.008 | 0.700 | 1.274 | 1.434 | 0.718 | 0.012 | >10 | 0.050 |

| >10 | 1-10 | 1-0.1 | 0.1-0.01 | <0.01 |
|---|---|---|---|---|
